# Transcriptional Condensates Encode a “Golden Mean” to Optimize Enhancer-Promoter Communication across Genomic Distances

**DOI:** 10.1101/2025.05.27.656519

**Authors:** Tao Zhu, Chunhe Li, Xiakun Chu

## Abstract

Enhancers regulate gene expression by physically contacting their target promoters, yet these contacts often span large genomic distances. Phase-separated condensates (droplet-like clusters) of transcription factors (TFs) are thought to facilitate such long-range enhancer-promoter (E-P) communication, but the quantitative principles underlying this mechanism remain unclear. Here we use polymer-based chromatin simulations to systematically vary the strength of TF clustering and the E-P genomic distance, examining their combined effects on E-P contact formation. We find that E-P contact frequency shows a non-monotonic dependence on the degree of TF clustering: contact frequency peaks at an intermediate TF abundance and TF-TF affinity, leading to a “golden mean” optimum. Two distinct regimes emerge: under weak TF-TF attraction, contact probability increases with condensate size, whereas strong TF attraction produces a peaked response that declines at high condensation levels. These results indicate that TF condensate acts as a tunable “rheostat”, buffering E-P interactions against increasing genomic distances. However, excessive TF clustering leads to molecular crowding and competition that ultimately impair E-P communication. Our study, consistent with recent experiments, establishes a mechanistic framework at molecular level, where balanced TF condensation enables robust long-range E-P communication, reconciling the stochastic nature of chromatin dynamics with the fidelity of gene regulation.

## Introduction

Enhancers play a central role in the mammalian genome by directing the spatiotemporal activation of genes through interactions with promoters, thereby ensuring precise control over developmental processes and cell-type-specific expression programs [1–4]. In general, enhancers can reside several kilobases to over a megabase away from their target promoters along the linear genome [5–8], productive transcription often requires that these distantly located elements come into close proximity and form physical contacts within the 3D space of the nucleus [9–13]. Yet how long-range enhancer-promoter (E-P) communication is accurately established and maintained remains poorly understood.

Chromosome conformation capture and imaging studies have demonstrated that E-P interactions occur predominantly within topologically associating domains (TADs), which are megabase-scale chromatin loops bounded by CTCF-cohesin complexes [14–17]. These structural units, characterized by frequent internal interactions, create a favorable environment for regulatory communication between distal genomic elements. Perturbation of TAD architecture can disrupt E-P contacts and lead to pathogenic gene misexpression [18–20]. However, transcriptional output has been often observed to remain only modestly affected following loop domain elimination through cohesin depletion [21–23]. This suggests that compensatory mechanisms may help stabilize gene expression programs, independent of the contribution of 3D genome structure. Even within intact TADs, quantitative studies have demonstrated that the transcriptional impact of enhancers does not follow a uniform pattern but instead depends strongly on their genomic distance from promoters [24, 25]. By systematically varying the E-P genomic distance (*L*_*E*-*P*_) within TADs and measuring the strength of gene expression, up to a fivefold difference was observed between *L*_*E*-*P*_ = 0 kb and 200 kb [24]. Noteworthy, certain long-range E-P interactions spanning over 1,000 kb have also been proven critical for normal gene expression, and their disruption can lead to dramatic transcriptional changes and severe disease phenotypes [7, 8], demonstrating that genomic distance alone cannot fully account for transcriptional regulation. Together, these findings underscore the need to further investigate the mechanisms that ensure robust gene regulation across large genomic distances.

Over the past decade, transcription factors (TFs) have been well-recognized as key regulators of gene expression not only through DNA binding, but also by compartmentalizing biochemical reactions into high-concentration hubs, referred to as phase-separated condensates and formed via multivalent interactions among themselves and co-factors [26–29]. Most TFs contain both DNA-binding domains (DBDs), which anchor them to specific genomic loci, and low-complexity domains (LCDs), which mediate interactions with other TFs and co-factors. These LCD-LCD interactions are functionally linked to transcriptional control and are considered promising targets for modulating gene activity [29, 30]. Owing to the inherent capacity to bind and cluster cooperatively, TFs are naturally equipped to promote E-P communication across large genomic distances. Indeed, a recent study revealed a weaker-than-expected dependence of E-P contact efficiency on linear genomic separation, which may be attributed to the organization of TF condensates [31]. However, to what extent TF phase separation modulates the sensitivity of E-P interactions with respect to genomic distance remains to be elucidated.

Although mounting evidence has confirmed the positive role of TF condensates in mediating E-P interactions and regulating gene activation [32–35], emerging studies have revealed that excessive TF phase separation can suppress transcriptional output [36–39]. This duality highlights the so-called “Goldilocks principle”, whereby the strength of TF condensation must be finely tuned to support optimal transcriptional regulation and an imbalance in this process may compromise gene expression dynamics and cellular function [36, 40]. Despite these insights, the quantitative relationship between the size of TF condensates and its influence on target E-P interactions remains poorly characterized. Moreover, while abnormally enhanced transcriptional condensates have been implicated in pathological conditions such as hormone-related disorders [37], the molecular basis for how excessive TF phase separation leads to transcriptional repression continues to be an open question.

Here, we develop a polymer-based chromatin model and perform molecular dynamics (MD) simulations to dissect how TF phase separation and E-P genomic distances jointly govern E-P communication. We find that E-P contact probability depends non-monotonically on both TF abundance and TF-TF interaction strength, with intermediate levels of condensation maximizing E-P contact probability across genomic distances. Increasing TF abundance and interaction strength progressively reduces the disparity in contact probability between near and far E-P pairs, ultimately eliminating it at high TF condensation levels. By quantifying TF clustering around E-P loci, we establish a link between condensate size and its influence on E-P contact formation, identifying two distinct regimes: under weak TF-TF interactions, E-P contact probability rises monotonically with condensate size, whereas under strong interactions, a non-monotonic “golden mean” relationship emerges. Finally, analysis based on the free energy landscape reveals that optimal phase separation reflects a balance between TF-mediated bridging that stabilizes the E-P complex and competitive TF occupancy that disrupts it. Together, these findings provide a mechanistic framework for understanding how finely tuned TF phase separation consistently helps ensure effective and efficient transcription across genomic distances.

## Results

### E-P communication with varying genomic distances and degrees of TF phase separation

We used a polymer-based chromatin model to investigate how different E-P genomic distances (*L*_*E*-*P*_), in conjunction with different degrees of TF phase separation, which are jointly modulated by the number of TF (*n*_TF_) and TF-TF interaction strength (*ϵ*_TF_), influence E-P communication. Our chromatin model adopts a beads-on-a-string representation, with each bead modeling a 30-nm chromatin fiber corresponding to a 3-kbp fragment of DNA [41, 42]. We considered a TAD of a typical genomic length 1,000 kb [14], within which we constructed six systems with different *L*_*E*-*P*_ values: 60, 200, 400, 600, 800 and 940 kb, respectively (Fig. 1A, left). Each system contains a single specific pair of enhancer and promoter loci. The two boundary ends of the TAD were connected and fixed in space (Fig. 1A, right), mimicking the structural constraints imposed by CTCF-cohesin complexes. Due to this structural setup, the spatial distance between the enhancer and promoter for *L*_*E*-*P*_ = 940 kb is comparable to that of *L*_*E*-*P*_ = − 60 kb. Thus, we denoted *L*_*E*-*P*_ = 940 kb as *L*_*E*-*P*_ = − 60 kb to emphasize their spatial similarity (600 and 800 kb were denoted as −400 and −200 kb, respectively). For simplicity, we modeled a single type of transcription factor that has the same fixed affinity with the enhancer/promoter and the variable affinity among themselves, while exhibiting purely volume-exclusion interactions with all other chromatin loci.

**Figure 1.**
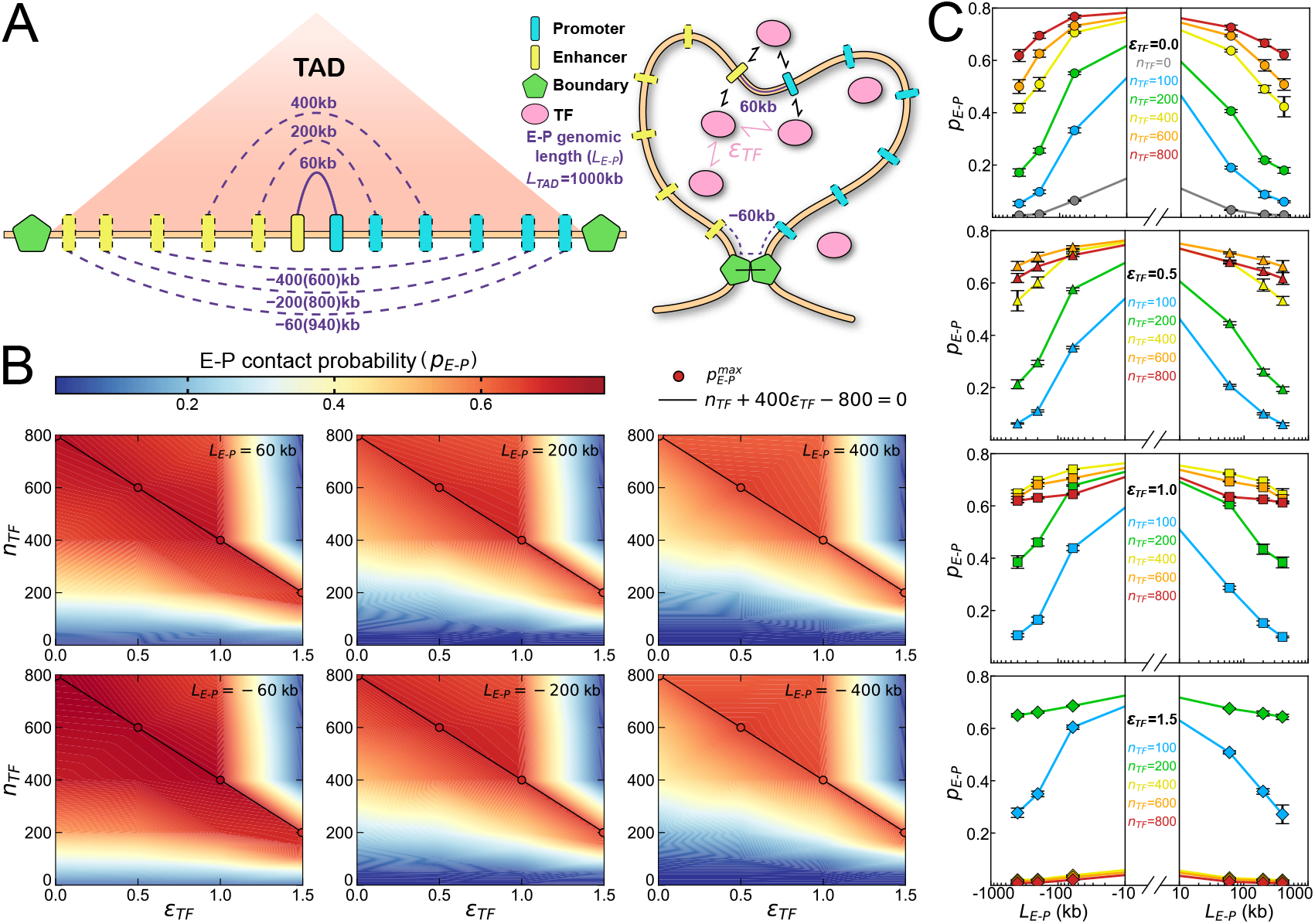
E-P contact patterns with various genomic distances and degrees of TF phase separation. (A) Schematic illustration of the polymer-based chromatin model, shown as a linear DNA-sequence configuration (left) and its spatial conformation (right), for a 1,000 kb TAD. The chromatin chain is colored brown and TAD boundaries are colored green, mimicking CTCF and cohesin proteins that exert structural constraints connecting two ends of TAD. The enhancer and promoter are each represented by a coarse-grained bead, colored yellow and turquoise, respectively. Six separate simulation systems, each featuring one of six distinct E-P genomic lengths (*L*_*E*-*P*_), are indicated by purple lines. Given the spatial similarity and symmetry, the 940, 800, and 600 kb systems are denoted as −60, −200, and −400 kb, corresponding to 60, 200, and 400 kb. Transcription factors (pink ovals) bind both enhancer and promoter with fixed affinity, while their multivalent interaction strengths are modulated by the parameter *ϵ*_TF_. (B) E-P contact probability (*p*_*E*-*P*_) maps for six different *L*_*E*-*P*_ systems with respect to the *ϵ*_TF_ and *n*_TF_, which collectively modulate the degree of TF phase separation. Red and blue regions represent high-contact and low-contact conditions, respectively. The red points 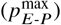 mark the positions whose values are the maximum in both their row and their column, and they all lie along the same straight line *n*_TF_ + 400*ϵ*_TF_ ™ 800 = 0 for all *L*_*E*-*P*_ systems. (C) An alternative representation of (B), divided into four panels for *ϵ*_TF_ = 0.0, 0.5, 1.0, 1.5, clearly showing the differences in *p*_*E*-*P*_ across different *L*_*E*-*P*_ settings and how these differences diminish as *n*_TF_ increases. Error bars represent the standard errors at the corresponding average values.

To quantitatively characterize the strength of E-P communication, we introduced the E-P contact probability [43, 44]:

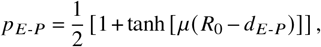

where *d*_*E*-*P*_ denotes the spatial distance between the enhancer and promoter, *R*_0_ is the cutoff distance defining contact, and *μ* controls the steepness of the function. We performed MD simulations for six different *L*_*E*-*P*_ systems, systematically varying both *ϵ*_TF_ and *n*_TF_ (details in Materials and Methods), and calculated the ensemble-average *p*_*E*-*P*_, as shown in Fig. 1B. Interestingly, we observed a consistent pattern of *p*_*E*-*P*_ across all values of *L*_*E*-*P*_: *p*_*E*-*P*_ does not monotonically vary with either *n*_TF_ or *ϵ*_TF_, or with both. Since there is a positive correlation between the tendency of TF phase separation and the number of TF and the strength of TF-TF binding affinity [45], this finding suggests that an optimal level of TF phase separation may possess a “golden mean” property for promoting E-P communication. Moreover, we found that when either *n*_TF_ or *ϵ*_TF_ was fixed, there existed a specific value of the other variable that maximized *p*_*E*-*P*_ for each *n*_TF_ = 200, 400, 600, 800 and *ϵ*_TF_ = 0.0, 0.5, 1.0, 1.5, respectively. Remarkably, these optimal combinations aligned along a linear relation: *n*_TF_ + 400*ϵ*_TF_ ™ 800 = 0, consistently across all six *L*_*E*-*P*_ values. This reveals a complementary relationship between TF concentration and TF-TF multivalent interaction strength in facilitating E-P contacts. Accumulating experimental evidence has shown that excessive TF phase separation or condensate formation can negatively impact gene expression [36–38], which aligns with the blue upper-right corners observed in all panels of Fig. 1B, indicating that excessive TF phase separation may impair E-P contact formation. Additionally, experimental studies have reported that transcriptional activation requires an optimal strength level of TF-TF multivalent interactions [36, 37], which is clearly captured in our contact maps: for example, at *L*_*E*-*P*_ = 200 kb and *n*_TF_ = 400, *p*_*E*-*P*_ exhibits a non-monotonic dependence on *ϵ*_TF_, reaching a maximum at *ϵ*_TF_ = 1.0. Another study revealed that increasing TF concentration or enhancing TF-TF multivalent interactions can have distinct effects on gene expression [38] : the former tends to maintain genes in a high-expression state, while the latter can suppress genes to a low-expression state. This observation is consistent with the trends observed along two distinct paths in our contact maps: for example, starting from the point *ϵ*_TF_, *n*_TF_ = (1.0, 400) in the panel of 200 kb, one path moves vertically to (1.0, 800), and the other horizontally to (1.5, 400). Notably, the mechanism by which excessive TF phase separation impairs gene activation remains unclear, and we will investigate this further in the following sections.

To better illustrate how *p*_*E*-*P*_ depends on *L*_*E*-*P*_, we presented the data in Fig. 1C. Two distinct *L*_*E*-*P*_-dependent modes of *p*_*E*-*P*_ were observed: one exhibits a rapid decline (e.g., the blue curves in the top panel), while the other shows a more gradual decrease (e.g., the red curves in the top panel). These modes are consistent with experimental observations [46], which attribute such differences to gene-specific features. Notably, our results suggested that TF condensation can modulate the extent to which *p*_*E*-*P*_ depends on *L*_*E*-*P*_. To more clearly evaluate this dependence, we selected 400 kb as a baseline and calculated the fold change in *p*_*E*-*P*_ for 60 kb and 200 kb relative to 400 kb under various combinations of *n*_TF_ and *ϵ*_TF_ (Fig. S1). We found that the presence of TFs eliminates the difference in *p*_*E*-*P*_ across genomic distances compared to the TF-free condition, consistent with experimental evidence [25]. For example, when *ϵ*_TF_ = 0 and *n*_TF_ = 100, the *p*_*E*-*P*_ value for *L*_*E*-*P*_ = 60 kb is more than threefold that of 400 kb. However, the disparity in *p*_*E*-*P*_ between 60 kb and 400 kb narrows with increasing *n*_TF_ and even becomes negligible at *ϵ*_TF_ = 1.0, 1.5 (Fig. 1C, S1). We concluded that sufficiently high degree of TF phase separation can compensate for the disadvantage of longer genomic distances in facilitating E-P contact. Interestingly, when *ϵ*_TF_ = 1.5 and *n*_TF_ ≥ 400, *p*_*E*-*P*_ drops uniformly across all *L*_*E*-*P*_ values, even falling below the baseline observed in the absence of TFs (the gray curves). Together, these findings reveal similar characteristics of E-P contact under varying *L*_*E*-*P*_ conditions, highlighting both the intrinsic influence of genomic distance and how this influence can be modulated by different degrees of TF phase separation.

### Quantifying the dependence of E-P contact on TF phase separation

To clearly recognize the size of TF condensates around the E-P loci, and how this influences *p*_*E*-*P*_, we calculated the number of TFs within the TF cluster enclosing the E-P pair (denoted as 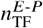, details in Materials and Methods) for various combinations of *n*_TF_, *ϵ*_TF_ and *L*_*E*-*P*_. We then plotted *p*_*E*-*P*_ as a function of 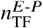 (Fig. 2A, B, S2). Interestingly, we observed two distinct trends depending on the strength of TF-TF multivalent interactions: in systems where these interactions were weak (Fig. 2A), the relationship was monotonic, whereas in systems with strong interactions (Fig. 2B), a non-monotonic “golden mean” behavior emerged. Transcription factors can often be categorized based on their functional roles into activators, repressors, and bifunctional regulators [47]. Notably, a recent study [48] reported a dual-role TF that can switch its transcriptional function depending on the expression level of its target gene, thereby recruiting distinct cofactors. In our case, the results suggest that increasing the intrinsic clustering propensity of TFs can fundamentally reshape how they regulate E-P contacts, with shifting from a purely promoting mode to a dual-mode involving both promotion and inhibition. This provides a perspective on how transcription factors may control gene expression through condensation-mediated mechanisms.

**Figure 2.**
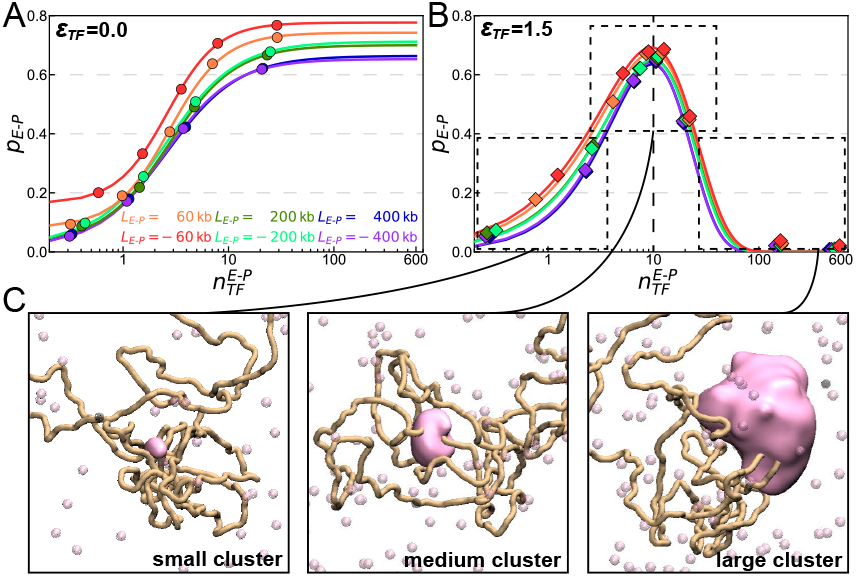
Quantitative analysis of E-P contact probability dependence on E-P-mediated TF condensate size. (A) *p*_*E*-*P*_ plotted against 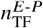 for six *L*_*E*-*P*_ settings when *ϵ*_TF_ = 0.0, where 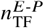 is the number of TFs in the cluster enclosing the E-P pair. Data points for each *L*_*E*-*P*_ are fitted using the Hill function. (B) Similar with (A) but with a non-monotonic trend, indicating a “golden mean” behavior emerges for *p*_*E*-*P*_ when TF-TF interaction is strong (*ϵ*_TF_ = 1.5). Data points for each *L*_*E*-*P*_ are fitted using a Generalized Gamma-like function. The results for the other two *ϵ*_TF_ values (0.5 and 1.0) are shown in Fig. S2. (C) Three representative snapshots from regions in (B): two low-*p*_*E*-*P*_ regions: one with insufficient and one with excessive E-P-mediated TF condensates; one high-*p*_*E*-*P*_ region with intermediate 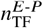. The snapshots highlight distinct sizes of E-P-mediated TF condensates (pink surfaces), defined as clusters where the maximum distance between any two TF beads is less than 3*σ* (*σ*: bead diameter). Color code is consistent with Fig. 1A.

Previous simulation and experimental studies revealed a sigmoid-like relationship between gene activation and TF concentration [49–51]. Inspired by these studies, we fitted our results using the Hill function respectively for different *L*_*E*-*P*_ settings. We observed that for the same TF condensate size, E-P pairs with shorter *L*_*E*-*P*_ exhibit higher contact probabilities. Conversely, to achieve the same *p*_*E*-*P*_, E-P pairs with shorter *L*_*E*-*P*_ require fewer TFs. These results reflect the dependence of E-P contact on genomic distance. In contrast, to our knowledge, no prior studies have reported trends similar to the non-monotonic pattern shown in Fig. 2B. We employed a Generalized Gamma-like function (details in Materials and Methods) to fit these data, which yielded good agreement. The results reveal a similar pattern across different *L*_*E*-*P*_ settings: there exists an optimal number of E-P-mediated TFs, approximately 10, that maximizes *p*_*E*-*P*_. Beyond this number, TFs begin to impair contact formation, consistent with a quantitative experimental result [50]. We concluded that the TF condensate surrounding the E-P loci must be maintained at a moderate size to most effectively promote E-P communication. To further illustrate this, we divided the data in Fig. 2B into three regions (Fig. 2C): two low *p*_*E*-*P*_ regions, one with insufficient and the other with excessive E-P-mediated TFs, one high *p*_*E*-*P*_ region with an intermediate 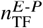. For each region, we presented representative snapshots from corresponding trajectories, which visually depict the size of the TF condensates around the E-P loci. These pictures clearly reflect the non-monotonic relationship between *p*_*E*-*P*_ and TF condensate size, consistent with experimental observations [39]. Together, our findings offer a more intuitive understanding of the conclusion from Fig. 1B, where TF phase separation possesses a “golden mean” property in facilitating E-P communication. In addition, once 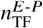 reaches or exceeds the optimal value, differences in *p*_*E*-*P*_ and 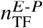 across varying *L*_*E*-*P*_ become significantly smaller compared to the regime before the optimum. This indicates that when TF condensates become excessively large, the dependence of E-P contact on genomic distance is markedly reduced. We speculate that under high degrees of TF phase separation, the search process for enhancer and promoter loci in 3D space is accelerated especially for the large *L*_*E*-*P*_ conditions. In other words, once a sufficiently large TF condensate forms, genomic distance no longer poses a substantial barrier to E-P communication.

### Genomic distance-dependence of E-P contact kinetics weakens as TF phase separation strengthens

To quantify the timescale of the search process of E-P pairs with different genomic distances, we calculated the mean first passage time (MFPT) under varying phase separation conditions (Fig. 3), where the contact threshold was defined as a spatial distance of 2*σ* (*σ* being the diameter of polymer beads). The MFPT resembles the initiation time of gene activation and the inter-locus relaxation time employed in chromosome-dynamics studies [31]. We observed that when *ϵ*_TF_ ≤ 1.0, increasing *n*_TF_ consistently reduces MFPT across all *L*_*E*-*P*_ settings. Likewise, at fixed *n*_TF_, the MFPT decreases with increasing *ϵ*_TF_. However, under conditions of strong TF-TF interactions (Fig. 3D), the MFPT exhibits a non-monotonic dependence on *n*_TF_. Specifically, at *n*_TF_ = 800, the MFPT for all *L*_*E*-*P*_ settings become even larger than those observed in the TF-free system (gray points in Fig. 3A). This indicates that excessive TF phase separation can paradoxically reduce E-P contact efficiency below that of passive 3D diffusion.

**Figure 3.**
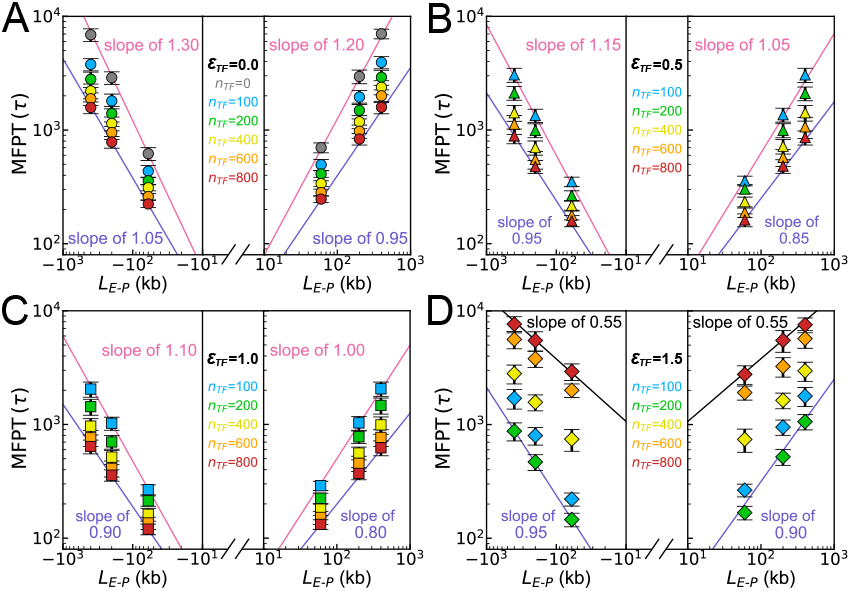
Kinetics of E-P contact formation and its dependence on genomic distance under various degrees of TF phase separation. (A-D) MFPT for the formation of E-P contact (E-P spatial distance ≤ 2*σ, σ*: bead diameter) from a spatially distant state, plotted against *L*_*E*-*P*_ on double-logarithmic axes. Data are grouped by *ϵ*_TF_ = 0.0, 0.5, 1.0, and 1.5 into four panels, with increasing *n*_TF_ in each and the color code consistent with Fig. 1C. Pink and purple lines delineate the range of data slopes, where slope magnitude reflects the MFPT dependence on *L*_*E*-*P*_. There is an exceptional case in (D), i.e., the black line (*ϵ*_TF_ = 1.5, *n*_TF_ = 800), where the system exhibited extreme TF phase separation, yielding significantly higher MFPT across all *L*_*E*-*P*_ values compared to (A-C), hence producing a slope that is not comparable to the pink line. *τ* is the reduced unit of time. Error bars represent the standard errors at the corresponding average values.

Increasing either *n*_TF_ or *ϵ*_TF_ reduced the steepness of the MFPT curves (Fig. S3), suggesting that the enhancement of TF condensates weakens the sensitivity of MFPT to *L*_*E*-*P*_. To quantify this effect, we presented the slope range of MFPT-*L*_*E*-*P*_ relationships under each *ϵ*_TF_ condition (Fig. 3, details in Materials and Methods). Given the symmetry between positive and negative *L*_*E*-*P*_ values, we observed an overall reduction in the MFPT slope from 1.30 to 0.80 upon enhancing TF phase separation (excluding an abnormal case: the black curve with the slope 0.55, see Materials and Methods for details). It should be noted that our model incorporated an explicit attractive interaction between E and P to represent their biochemical compatibility and confined them within a loop domain, facilitating their contact [52]. Accordingly, the maximal slope of 1.30 falls below the theoretical exponent of 2 predicted by the simple Rouse polymer model. Intriguingly, the minimal slope, 0.80, deviates further from 2 and is close to a recent experimental measurement of 0.7 ± 0.2 reported in a landmark study [31], supporting that E-P pairs separated by large genomic distances still have a considerable chance of forming specific contacts. Previously, the discrepancy between experimental measurements and the ideal Rouse polymer model was attributed to the heterogeneities of E-P pairs, regulated by a number of biological process including the TF clustering [31]. We emphasized that by simply modulating the intensity of TF condensates, the slope characterizing the transcriptional dependence on genomic distance, can be reduced to values very close to experimental measurements, shedding light on the potent role of TF condensates in orchestrating genome structural dynamics and function.

### Thermodynamic and molecular insights into the bidirectional modulation of gene activation by TF phase separation

To investigate the underlying mechanism by which different degrees of TF phase separation differ significantly in mediating E-P communication, we defined a reaction coordinate *ξ* = *n*_TF_ + 400*ϵ*_TF_ ™ 800, which integrates TF number and TF-TF binding affinity into a single measure of TF condensation intensity. As shown in Fig. 1B, *p*_*E*-*P*_ can be maximized at *ξ* = 0. We then quantified the 2D free energy landscape of E-P communication as a function of *p*_*E*-*P*_ and the E-P spatial distance (*d*_*E*-*P*_), with the formula:

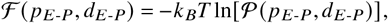

where 𝒫 (*p*_*E*-*P*_, *d*_*E*-*P*_) denotes the joint probability distribution of chromatin conformation at specific values *p*_*E*-*P*_ and *d*_*E*-*P*_ (details in SI Appendix). Since *p*_*E*-*P*_ exhibits similar TF-modulated patterns across all *L*_*E*-*P*_ settings (Fig. 1B), we fixed *L*_*E*-*P*_ = 400 kb and then presented three energy landscapes at *ξ* = − 400, 0, 400, respectively (Fig. 4A). Each landscape revealed three stable states consistent with our previous work [51] and experimental evidence [53], corresponding to a two-step contact-formation pathway: transitions from O_off_ to P_off_, and finally to P_on_. The three states were defined as: the P_on_ state, where E-P contact is established and ready for gene activation; the P_off_ state, where the E-P pair is in spatial proximity but not close enough to establish contact, thus may not trigger transcription; and the O_off_ state, where the enhancer and promoter are spatially far from each other. With increasing *ξ*, the global stability of P_off_ increases and that of O_off_ decreases monotonically, while P_on_ exhibits a non-monotonic trend that peaks at *ξ* = 0.

**Figure 4.**
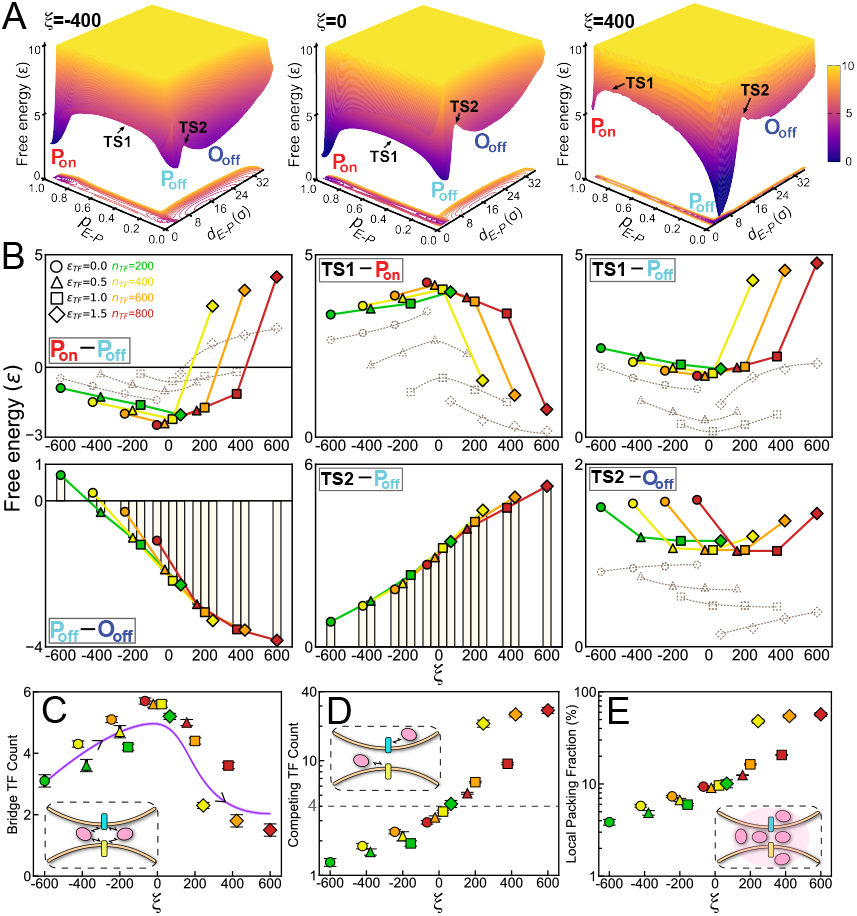
Thermodynamic analysis and molecular-mechanistic quantification of E-P contact mediated by TF condensation intensity. (A) 2D free energy landscape projected onto *p*_*E*-*P*_ and *d*_*E*-*P*_ with respect to TF condensation intensity *ξ*, when *L*_*E*-*P*_ = 400 kb. Three stable states are observed: the P_on_ state, where the enhancer and promoter are spatially close and form a contact; the P_off_ state, where they are spatially close but not enough to form a contact; and the O_off_ state, where they are spatially distant without contact. The transition states between P_on_ and P_off_, and between P_off_ and O_off_, are indicated as TS1 and TS2, respectively. (B) Free energy differences between the stable states and the barrier heights during the state-transitions as a function of *ξ*. Solid colored markers and lines indicate non-monotonic trends observed when varying *ϵ*_TF_ at fixed *n*_TF_, while dashed gray markers and lines represent non-monotonic trends observed when varying *n*_TF_ at fixed *ϵ*_TF_. (C-E) Three quantities are defined to explain the microscopic mechanism by which TFs mediate E-P contact, illustrated by inset figures: the bridge TF count, which is the number of TFs simultaneously in close interaction with both enhancer and promoter; the competing TF count, which is the number of TFs in close interaction with either enhancer or promoter but not both; and the local packing fraction, which is the TF occupancy at the E-P loci.

We then calculated the free energy differences as a function of the reaction coordinate *ξ*, including (i) those between stable states (reflecting relative stability) and (ii) those between stable and transition states (TSs) (reflecting barrier heights) (Fig. 4B), where *ξ* includes 16 combinations of *ϵ*_TF_ = 0.0, 0.5, 1.0, 1.5 and *n*_TF_ = 200, 400, 600, 800, corresponding to seven distinct degrees of phase separation. Interestingly, increasing *ξ* initially stabilizes the P_on_ state relative to the P_off_ state, but further enhancement leads to a reversal in stability. Correspondingly, the transition from P_off_ to P_on_ is first accelerated, then substantially inhibited. Regarding the unbinding dynamics of E-P contact, a non-monotonic trend was also observed, with the optimal *ξ* = 0 consistent with the two former trends. In contrast, *ξ* has little effect on the first transition step from the O_off_ state to the P_off_ state, while a subtle non-monotonic trend emerges, of which the rising phase recalling the observations in Fig. 3D. Moreover, the stability of the P_off_ state relative to the O_off_ state monotonically increases with increasing *ξ*, mainly due to the increase in the barrier height between TS2 and P_off_ when the TF condensation intensity is enhanced.

To reveal the mechanism underlying the non-monotonic trends among P_on_, P_off_, and TS1, we defined and calculated three quantities (Fig. 4C, D, E; SI Appendix): the bridge TF count (BTC), defined as the number of TFs simultaneously in close interaction with both enhancer and promoter, which mediate the bridge effect; the competing TF count (CTC), defined as the number of TFs in close interaction with either enhancer or promoter but not both, which mediate competitive binding effect; and the local packing fraction (LPF), defined as the TF occupancy at the E-P locus. Intuitively, BTC initially increases with increasing *ξ* due to higher TF abundance and affinity that favor establishment and maintenance of the P_on_ state. This corresponds to the initial decline in the TS1-P_off_ barrier and rise in the TS1-P_on_ barrier. Notably, for *ξ* > 0, the LPF becomes pronounced, causing BTC to decline as overcrowded TFs weaken their own bridging effect for E-P contact. Concurrently, CTC surpasses BTC and the gap widens, reflecting an increasingly dominant competitive binding effect. Together, these effects elevate the TS1-P_off_ barrier and lower the TS1-P_on_ barrier. In conclusion, our molecular-mechanistic analysis demonstrated that TFs exert both bridge and competing effects on E-P communication under varying TF condensation intensities; the relative strengths of these effects change with *ξ*, collectively producing the observed “golden mean” behavior in the relative stability of the P_on_ state and in its global stability (*p*_*E*-*P*_), thereby illuminating the intricate mechanism by which TF condensates bidirectionally modulate gene activation.

## Discussion and Conclusions

Taken together, we have demonstrated that a “golden mean” modulation governed by TF phase separation consistently optimizes E-P communication efficiency across varying genomic distances (Fig. 5). From a thermodynamic perspective, optimal transcriptional efficiency requires a moderate proportion of the P_off_ state (spatially proximal but transcriptionally inactive) and a minimal presence of the O_off_ state (spatially distant) (Fig. 5A, B, S4). As TF condensation enhances, the P_off_ state becomes increasingly stabilized. We propose that this stabilization, within an appropriate range of TF phase separation, reflects TF accumulation around the E-P loci, thereby facilitating transitions to the active P_on_ state and extending its lifetime. This interpretation echoes recent experimental observations showing that an intermediate E-P contact stage, where numerous proteins and RNAs accumulate around the gene, can extend the duration of subsequent transcription activities [54]. Notably, we have found that excessive TF condensation over-stabilizes the intermediate P_off_ state, hindering transitions to the active P_on_ state and thereby reducing transcriptional efficiency. In this context, maintaining the P_off_ state at a balanced, “golden mean” level appears essential for efficient transcriptional communication. Meanwhile, the near-zero percentage of the O_off_ state indicates that enhancers and promoters are almost always in close proximity, suggesting that the entropic constraints imposed by genomic distance can be effectively overcome by sufficiently strong TF condensates.

**Figure 5.**
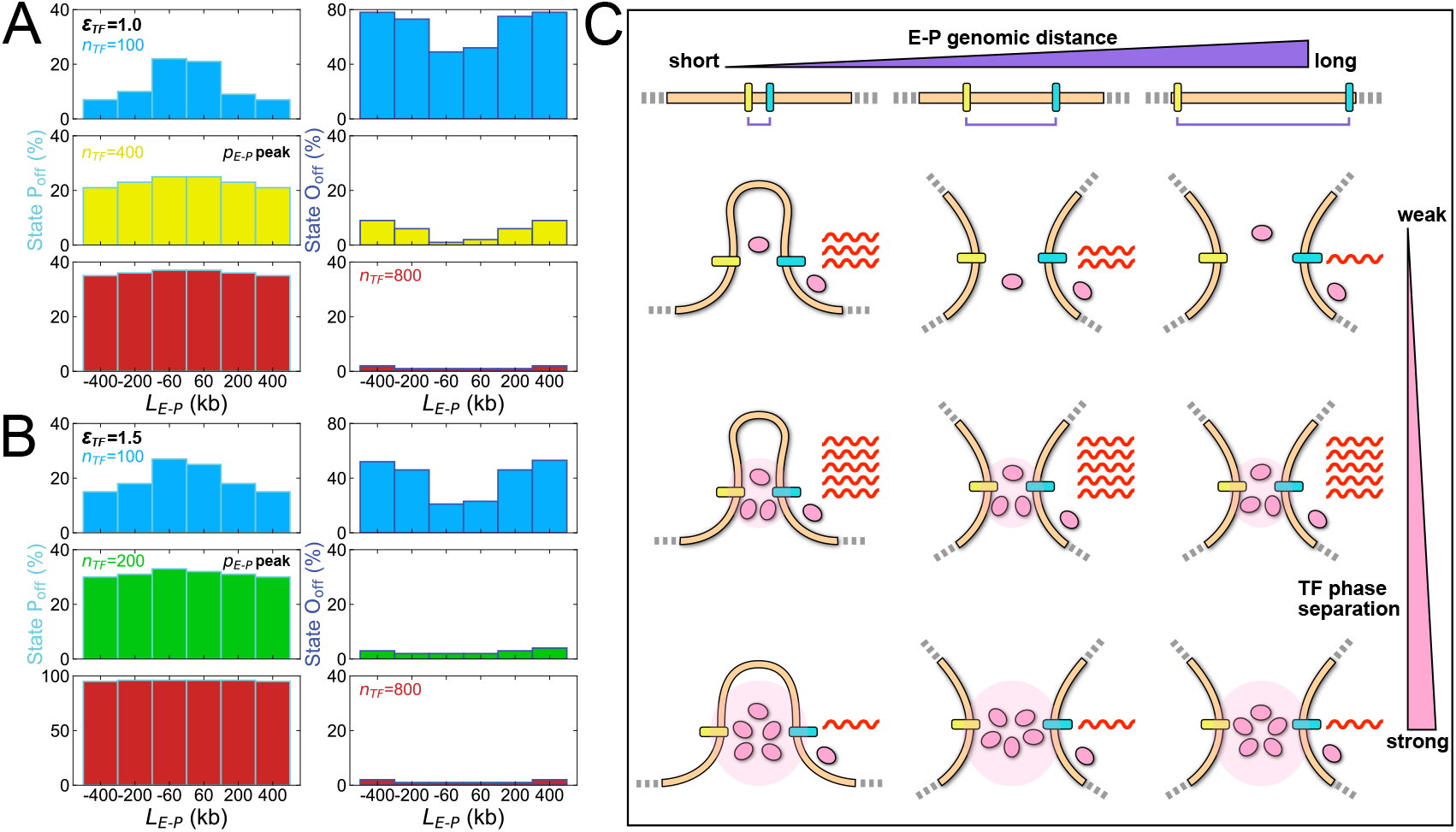
“Golden mean” of TF phase separation for efficient E-P communication across all genomic distances. (A-B) Global stability (percentage) of the P_off_ and O_off_ states for E-P contact under representative TF conditions. The optimal ranges to achieve the highest *p*_*E*-*P*_ are 20-30% for P_off_ and 0-10% for O_off_ across all *L*_*E*-*P*_ values (yellow and green). When the O_off_ percentage drops below 10% and approaches 0%, the differences in *p*_*E*-*P*_ across different *L*_*E*-*P*_ settings become negligible (see Fig. 1C). The results for the other two ϵ_TF_ values (0.0 and 0.5) are shown in Fig. S4. (C) Schematic illustration summarizing the main findings of the study.

The comparable E-P contact probabilities observed at *L*_*E*-*P*_ = ±60 kb recall the critical role of the CTCF-cohesin complex in organizing long-range E-P interactions [55–57]. In contrast, the similar *p*_*E*-*P*_ values observed at *L*_*E*-*P*_ = −60 kb and −400 kb under strong condensation conditions mirror the experimental observations that condensate-driven chromatin looping can occur independently of CTCF-cohesin function [58]. These observations reinforce the emerging model in which transcriptional condensates dynamically tether enhancers and promoters to confer spatial specificity [59, 60]. Moreover, the minimum slope of 0.80 observed for MFPT as a function of *L*_*E*-*P*_ (Fig. 3) agrees well with the experimental measurement of 0.7 ± 0.2 [31], highlighting a weaker-than-expected dependence of contact kinetics on genomic separation when modulated by TF condensation. This observation may help explain how certain developmental genes, such as *Myc*, maintain robust transcriptional activity despite megabase-scale separations [7, 8, 61]. More broadly, our work has established a theoretical framework for understanding transcriptional condensation as a tunable “rheostat” that enables regulatory elements to overcome spatial constraints, reconciling the stochastic nature of chromatin dynamics with the precision demands of developmental transcription.

Although gene regulation by TF condensates is well established as essential for facilitating differentiation, driving cell state transitions, and maintaining cellular functions [62–64], emerging evidence suggests that abnormal TF condensation can impair transcription [36–39]. Here, we have quantitatively revealed the relationship between the size of transcriptional condensates and the efficiency of target E-P contacts (Fig. 2). Interestingly, under two distinct regimes of weak and strong TF-TF interactions, the regulation of E-P communication by TF condensates exhibits two functional patterns: a sigmoidal response and a “golden mean” behavior. By simply increasing the strength of TF-TF interactions, we obtained two functional types of TFs: one that monotonically enhances E-P contact probability, and another that regulates it bidirectionally depending on the degree of condensation. These findings may inform strategies targeting TF-TF interactions for modulating transcriptional output [29, 65]. Mechanistically, we further demonstrate that excessive TF condensation inhibits E-P contact by reducing the stability of the transcriptionally active state, P_on_. To elucidate this, we classify TFs surrounding the E-P loci into two functional groups: one group associates with both enhancer and promoter simultaneously, functioning like “bridging” proteins that have been described to facilitate genome reorganization [42, 66, 67]; the other group binds exclusively to either the enhancer or promoter, effectively competing for available binding sites and thereby interfering with E-P association. This competitive effect is reminiscent of biochemical titration mechanisms observed in live cells [68–70]. As TF condensation increases, the size of the bridging TF group displays a non-monotonic trend that mirrors the relative stability of P_on_ compared to P_off_. In contrast, the competing TF group expands monotonically, exerting increasing inhibitory effects on both the association and dissociation of the E-P complex. The interplay between these two TF populations gives rise to the observed “golden mean” behavior in both the global and relative stability of the productive state P_on_, emphasizing the critical balance between bridging and competing TF functions required for optimal E-P communication (Fig. 5C).

Notably, numerous experimental and simulation studies have shown that TF condensates tend to solidify and lose liquidity under conditions of excessive TF-TF intersactions [37, 39, 45, 71, 72], which greatly reduces their capacity for internal reorganization. We speculate that this phenomenon may also occur in our system, where a jamming effect, together with a dominant competing mechanism, jointly contributes to the inhibition of E-P communication. An additional factor that may explain the increased energy barrier for transitions from the O_off_ to P_off_ state (Fig. 4B) is the emerging ability of TFs to self-condense without requiring mediation by E-P loci. Analysis of simulation snapshots reveals that TFs can indeed form condensates independently at high interaction strengths (ϵ_TF_ = 1.5) and TF numbers (*n*_TF_ = 400, 600, 800; Fig. S5). Such behaviors may lead to contact efficiencies resembling those of passive diffusion of E-P loci, aligning with experimental findings that report impaired target search by condensates when self-assembly becomes dominant [73]. From a physical perspective, we highlight the role of transcriptional condensation as a tunable “rheostat” capable of scaling inistial gene activation efficiency across genomic distances. However, our findings also underscore that this “rheostat” must operate within an optimal range because excessive condensation introduces structural rigidity and competition, which can hinder effective regulation. Our study offers both mechanistic insight and theoretical prediction, suggesting that TF condensates are flexible and versatile targets for regulating transcription, but the intensity of the condensation must be carefully modulated to avoid dysfunctional states.

## Materials and Methods

### Coarse-grained chromatin model and MD simulations

We coarse-grained chromatin into a polymer model using a beads-on-a-string representation, where each bead corresponds to a 30-nm chromatin fiber and spans approximately 3 kbp of DNA [41, 42]. We focused on a single TAD with a typical length of 1,000 kbp [14], corresponding to 333 consecutive beads on a polymer string. To better represent the flanking TAD context, 100 beads were appended to each end of the string beyond the TAD boundaries. The constraining effect of CTCF and cohesin proteins at TAD boundaries was implemented as a harmonic potential. Bonded interactions between adjacent beads (*i*,*i* + 1) were described by a harmonic potential:

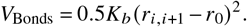

The stiffness of the chromatin chain was realized by a linear-placement potential among three adjacent beads (*i* − 1,*i*,*i* + 1):

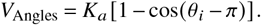

Non-bonded interactions between any two beads (*i, j*) that satisfy |*i* −*j*| > 2 were modeled using a Lennard-Jones potential:

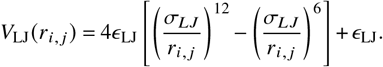

To mimic the confined TAD environment, we applied a spherical confinement potential with radius *R*_C_:

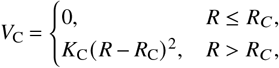

We then constructed six separate systems, each containing a single enhancer-promoter pair symmetrically positioned on either side of the TAD midpoint, with genomic separations of *L*_*E*-*P*_ = 60, 200, 400, 600, 800 and 940 kb, where each enhancer or promoter was represented by a single bead. For all systems, we introduced TFs modeled as free beads of the same size, diffusing around the chromatin chain. These TFs interact specifically with each other as well as with enhancers and promoters via Lennard-Jones potentials (*V*_LJ_), while exhibiting purely repulsive (volume-excluding) interactions with all other chromatin sites. Both the number of TFs and the strength of TF-TF interactions were varied to modulate different degrees of TF phase separation, with *n*_TF_ = 0, 50, 100, 150, 200, 400, 600, 800 and ϵ_TF_ = 0.5, 1.0, 1.5, where ϵ_TF_ is the strength parameter placed in front of *V*_LJ_. Noteworthy, when ϵ_TF_ = 0.0, only repulsive interactions between TFs were considered. Detailed setting of the parameters in the abovementioned potentials can be in SI Appendix.

All MD simulations were conducted using Gromacs (version 4.5.7) [74], with the Plumed plugin (version 2.5.0) [75] to implement spherical confinement. Simulations were performed in reduced units. Langevin dynamics was applied with a friction coefficient of 1.0*τ*^−1^, where *τ* denotes the reduced time unit. The integration time step was set to 0.0005*τ*, and the simulation temperature was fixed at ϵ, which is the reduced energy unit. The polymer bead diameter *σ* was used as the unit of length. For each combination of parameters (*L*_*E*-*P*_, ϵ_TF_, *n*_TF_), we performed 30 independent simulations initialized from distinct configurations. Each simulation was run for 40,000*τ*, with data collected from the final 20,000*τ* to ensure stable and sufficient sampling for downstream analysis.

### Quantifying E-P-mediated TF condensate size and its relationship with p_E-P_

We employed the Density-Based Spatial Clustering of Applications with Noise (DBSCAN) algorithm to quantify the size of E-P-mediated TF condensates [76], denoted as 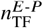. DBSCAN identifies clusters as groups of densely packed points, where each core point must have at least *N*_min_ neighbors within a radius *δ*. The algorithm expands clusters by connecting core points and their reachable neighbors. In our simulations, DBSCAN was applied to the combined set of all TF beads along with the enhancer and promoter beads in each frame. We set *N*_min_ = 5 and *δ* = 3*σ*, which represent the minimum cluster size and the maximum distance between two beads to be considered within the same cluster, respectively. For each frame, we identified the cluster (if any) that simultaneously contained both E and P. If such a cluster was found, the number of TFs within it was recorded as 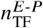; otherwise, 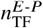 was set to zero. The final value was obtained by averaging 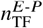across all frames and plotted as a function of 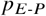.

To examine how *p*_*E*-*P*_ varies with 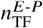, we fitted the data for each *L*_*E*-*P*_ value using two different functions for ϵ_TF_ = 0.0 and 1.5, respectively. For ϵ_TF_ = 0.0, the data exhibited a sigmoidal trend and was fitted using the Hill function, consistent with previous studies [49–51]:

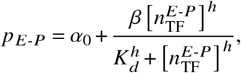

where four parameters were fitted: *h*, the Hill coefficient, *K_d_*, the effective binding affinity, and *α*_0_ and *β*, which determine the minimum and maximum values of *p*_*E*-*P*_, respectively. For ϵ_TF_ = 1.5, the data displayed a previously unreported non-monotonic trend, to the best of our knowledge, and was fitted using a Generalized Gamma-like function:

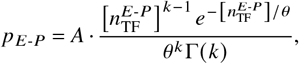

where *A* is a scaling factor, *k* is the shape parameter, *θ* is the scale parameter, and Γ(*k*) denotes the gamma function, a continuous extension of the factorial.

### Calculating MFPT and examining its exponential relationship with L_E-P_

To determine the MFPT for E-P contact formation, we defined the formation as *d*_*E*-*P*_ ≤ 2*σ*, corresponding to an instantaneous condition of *p*_*E*-*P*_ ≥ 0.98. It should be noted that all simulations performed earlier started from distinct initial configurations with *d*_*E*-*P*_ ≥ 10*σ* to ensure a spatially separated state. Therefore, we recorded the time required for E-P pairs to reach the formation threshold for the first time in simulations of each parameter set (*L*_*E*-*P*_, ϵ_TF_, *n*_TF_). The MFPT was then obtained by averaging the first passage time across 30 independent trajectories.

To characterize how MFPT depends on *L*_*E*-*P*_ and its modulation by TF phase separation, we adopted an exponential scaling relationship inspired by previous work [31]:

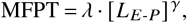

where *λ* denotes the overall timescale, and *γ* describes the dependence of MFPT on *L*_*E*-*P*_, corresponding to the slope in double-logarithmic axes. Given slight differences in timescales between positive and negative *L*_*E*-*P*_ data, we fixed *λ* = 5 and 3 for the two cases, respectively, and fitted *γ* for each combination of parameters (ϵ_TF_, *n*_TF_). We then reported the indicative minimum and maximum values of *γ* at each ϵ_TF_ panel to illustrate the effect of varying *n*_TF_. Notably, under the condition ϵ_TF_ = 1.5 and *n*_TF_ = 800, the system exhibited excessive TF phase separation, leading to consistently elevated MFPT values across all *L*_*E*-*P*_ settings. An appropriate *λ* for this case two orders of magnitude higher than that of other indicative lines (pink and purple lines in Fig. 3). Although the fitted *γ* = 0.55 (black lines) represents the indicative extremum, the significantly different timescale highlights the abnormal nature of this case, so this *γ* value should not be directly compared with the others. On the other hand, this minimal *γ* indicates that in a system with excessive TF phase separation, MFPT is least affected by *L*_E-*P*_, while E-P contact remains uniformly difficult across all genomic distances compared to systems with milder TF phase separation.

## Data Availability

The necessary files for setting up the Gromacs simulations with PLUMED and the analysis scripts are publicly available at GitHub (https://github.com/icecolaTao/ChrTFModel). It includes the following components: (1) simulation files: Gromacs input files for running the chromatin polymer model simulations; (2) analysis tools: Python scripts for downstream analysis of simulation output files, including calculations of TF clustering, MFPT and landscape. All other data are included in the manuscript and/or supporting information.

## Acknowledgments

X.C. was supported by the National Natural Science Foundation of China (grant No. 12474201 and No. 32201020), the general program of Guangdong Basic and Applied Basic Research Foundation (grant No. 2024A1515010862), the Guangdong Scientific Research Platform and Projects for Higher-educational Institutions from the Department of Education of Guangdong Province (grant No. 2023KTSCX169), and the Guangdong Provincial Project (grant No. 2023QN10X037). C.L. was supported by the National Natural Science Foundation of China (Grant No. 12171102), and the National Key R&D Program of China (Grant No. 2019YFA0709502). The authors also acknowledge the Green e Materials Laboratory (GeM) and HPC+AI Intelligence Computing Center at the Hong Kong University of Science and Technology (Guangzhou) for providing computational support.

## Notes

### Competing Interest Statement

The authors have declared no competing interest.

