## Supplementary material for "Transcriptional Condensates Encode a “Golden Mean” to Optimize Enhancer-Promoter Communication across Genomic Distances": SI Text

***"This document provides supporting text and supplementary figures (Figures S1-S8)."***

### Supporting text

#### Details on model and simulation parameters

We coarse-grained chromatin into a polymer model using a beads-on-a-string representation, where each bead corresponds to a 30-nm chromatin fiber and spans approximately 3 kbp of DNA [1, 8]. We focused on a single TAD with a typical length of 1,000 kbp [7], corresponding to 333 consecutive beads on a polymer string. To better represent the flanking TAD context, 100 beads were appended to each end of the string beyond the TAD boundaries. The constraining effect of CTCF and cohesin proteins at TAD boundaries was implemented as a harmonic potential. Bonded interactions between adjacent beads $(i, i+1)$ were described by a harmonic potential:

$$V_{\text{Bonds}} = 0.5K_b(r_{i,i+1} - r_0)^2.$$

The stiffness of the chromatin chain was favored by a linear-placement potential among three adjacent beads $(i-1, i, i+1)$:

$$V_{\text{Angles}} = K_a[1 - \cos(\theta_i - \pi)].$$

Non-bonded interactions between any two beads $(i, j)$ that satisfying $j - i > 2$ were modeled using a Lennard-Jones potential:

$$V_{\text{LJ}}(r_{i,j}) = 4\epsilon_{\text{LJ}}\left[\left(\frac{\sigma_{LJ}}{r_{i,j}}\right)^{12} - \left(\frac{\sigma_{LJ}}{r_{i,j}}\right)^{6}\right] + \epsilon_{\text{LJ}}.$$

To mimic the confined TAD environment, we applied a spherical confinement potential with radius $R_{\text{C}}$:

$$V_{\text{C}} = \begin{cases} 0, & R \leq R_C, \\ K_{\text{C}}(R - R_{\text{C}})^2, & R > R_C, \end{cases}$$

Simulations were performed in reduced units. $\tau$ denotes the reduced time unit. $\epsilon$ is the reduced energy unit. The polymer bead diameter $\sigma$ was used as the unit of length. The bonded potential was set with $r_0 = \sigma$ and $K_b = 3200\epsilon/\sigma^2$, preventing excessive extension or overlap between adjacent beads. The angular potential was applied with $K_a = 2.0\epsilon$ to maintain the chain stiffness. The non-bonded LJ potential within the TAD was set with $\sigma_{LJ} = 1.5\sigma$ and a uniform interaction strength $\epsilon_{LJ} = 0.15\epsilon$, allowing moderate chromatin compaction. Enhancer-promoter interactions used a stronger LJ strength $\epsilon_{LJ} = 2.0\epsilon$ to reflect their biochemical compatibility. TFs interacted with the enhancer and promoter with $\epsilon_{LJ} = 5.0\epsilon$, consistent with previous studies [10]. Volume-excluding interactions between TFs and other chromatin loci were represented by the repulsive part of the LJ potential $4\epsilon\left(\sigma/r_{i,j}\right)^{12}$. All non-bonded interactions were truncated at a cutoff distance of $7.5\sigma$. The radius of the spherical confinement was set to $R_{\text{C}} = 32\sigma$, with a strength of $K_{\text{C}} = 100\epsilon/\sigma^2$. For calculating the E-P contact probability ($p_{E\text{-}P}$), we used the parameters $\mu = 10.0\sigma$ and $R_0 = 2.5\sigma$. Values of $\mu$ and $R_0$ vary across studies [6, 9], suggesting limited sensitivity of the results to their specific choices.

#### Calculations of thermodynamic and molecular quantities of E-P communication

We quantified the 2D free energy landscape of E-P communication using the formula:

$$\mathcal{F}(p_{E\text{-}P}, d_{E\text{-}P}) = -k_B T \ln[\mathcal{P}(p_{E\text{-}P}, d_{E\text{-}P})],$$

where $\mathcal{P}(p_{E\text{-}P}, d_{E\text{-}P})$ denotes the joint probability distribution of chromatin conformations sampled across all trajectories. The variable $d_{E\text{-}P}$ corresponds to the spatial distance between the enhancer and promoter, while $p_{E\text{-}P}$, derived from $d_{E\text{-}P}$ via a nonlinear transformation, primarily characterizes the interaction process when the enhancer and promoter are in close proximity. Previous studies have shown that $d_{E\text{-}P}$ is well-suited for capturing unbound states where no contact forms, whereas $p_{E\text{-}P}$ provides a more sensitive description of the contact formation process [4, 5]. We then calculated the relative stability between the three stable states ($P_{on}$, $P_{off}$, $O_{off}$) and the barrier heights for transitions crossing the transition states TS1 and TS2 at seven $\xi$ values. Since direct calculations using the 2D landscapes can be cumbersome, we simplified the analysis by constructing a 1D landscape using a single reaction coordinate $\zeta_{E\text{-}P} = d_{E\text{-}P} - 5 \times p_{E\text{-}P}$ (Fig. S6), following our previous work [10].

To elucidate the non-monotonic energy differences among $P_{on}$, $P_{off}$, and TS1, We quantified three relevant quantities under conditions where the enhancer and promoter are in close proximity, defined as $d_{E\text{-}P} \leq K$, with $K$ representing the maximum distance at which the E-P pair remains in either the $P_{on}$ or $P_{off}$ state for each specific combination of $(\epsilon_{TF}, n_{TF})$. Bridge TF count (BTC) refers to the number of TFs simultaneously located within $2\sigma$ of both the enhancer and promoter, indicating TFs that are tightly associated with both elements and likely serve as molecular bridges. Competing TF count (CTC) captures the number of TFs that are within $2\sigma$ of either the enhancer or the promoter, but not both, and thus may compete for binding without contributing to E-P contact. Local packing fraction (LPF) is defined by centering a sphere of radius $5\sigma$ at the midpoint between the enhancer and promoter, then counting the number of TF beads inside this sphere, denoted as $n_{TF}^{sphere}$. The LPF is calculated as the total volume occupied by TF beads divided by the sphere's volume. However, due to volume exclusion effects, TFs cannot fully fill the sphere. We therefore determined the maximal number of TFs that can fit inside such a sphere under stronger TF-TF attraction ($\epsilon_{TF} = 2.0$) to be 109, and used this value for normalization, yielding LPF = $100\% \times n_{TF}^{sphere}/109$.

We further presented results analogous to those in Fig. 4, but for $L_{E\text{-}P} = 200$ kb (Figs. S6, S7 and S8), demonstrating the robustness of our findings. The main differences compared to the $L_{E\text{-}P} = 400$ kb case lie in the facilitated transition from the $O_{off}$ to $P_{off}$ state and the consequently enhanced stability of the $P_{off}$ state at $L_{E\text{-}P} = 200$ kb. Notably, when $L_{E\text{-}P} = 60$ kb, $\epsilon_{TF} = 1.5$, and $n_{TF} \geq 400$ (Fig. S8), the $O_{off}$ state vanished due to strong TF phase separation, which promotes the enhancer and promoter to reach and remain in close spatial proximity, coupled with the inherently short genomic distance.

#### Estimated correspondence from reduced units to physical units

In our simulations, we define the unit of length $\sigma$, as the diameter of each bead, which represents a 30-nm chromatin fiber. This chromatin fiber has been reported to hold a persistence length between 170 and 220 nm [3], which we used to estimate the physical scale of $\sigma$. A recent live-cell imaging study in *Drosophila* investigated the motion of an enhancer-promoter pair separated by 58 kb of DNA [2], a length comparable to the 60 kb separation used in our simulations. This study reported that the mean squared displacement (MSD) of the promoter reached approximately 900,000 $nm^2$ at 200 seconds. By comparing our simulation results with this experimental observation, we estimated our time unit $\tau$ to lie between 1.7 and 2.9 seconds.

### Supplementary figures

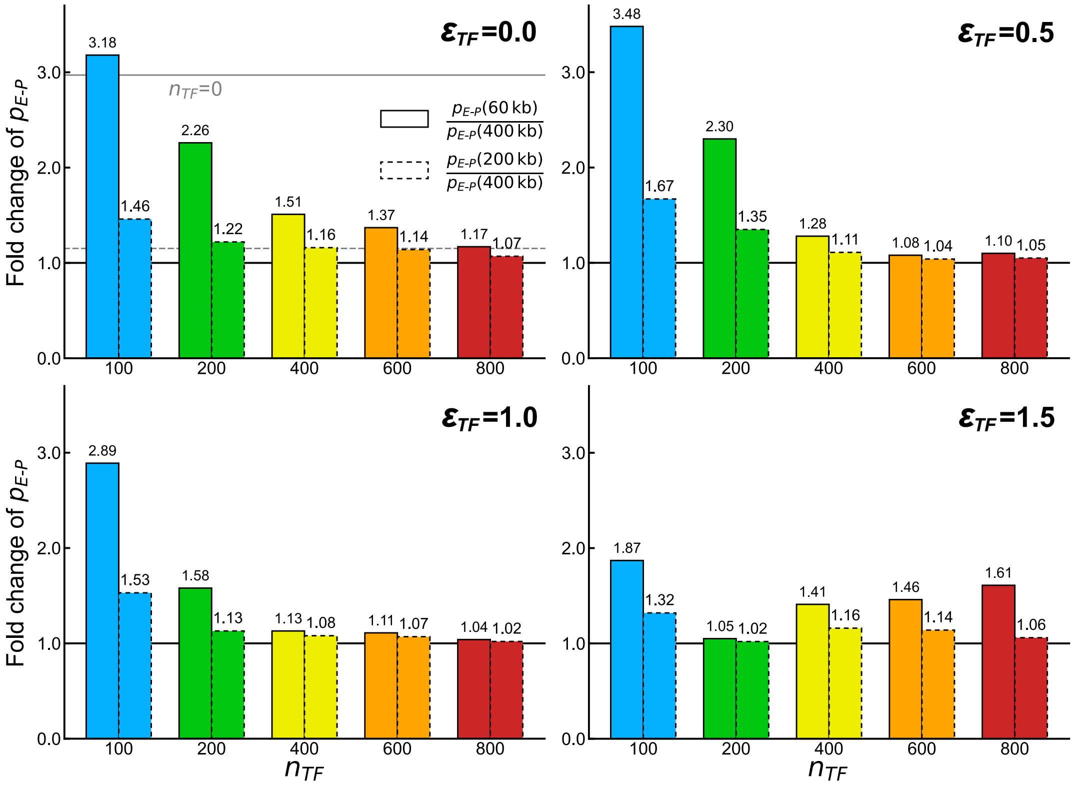


**Figure S1:** Fold change in E-P contact probability for $L_{E\text{-}P}$ = 60 and 200 kb relative to that of $L_{E\text{-}P}$ = 400 kb, as $n_{\mathrm{TF}}$ increases at each $\epsilon_{\mathrm{TF}}$ value. The $p_{E\text{-}P}$ values correspond to those shown in Fig. 1C in the main text. A fold change value closer to 1 indicates that E-P contact becomes less dependent on E-P genomic distance, suggesting the effect of modulation by TF phase separation.

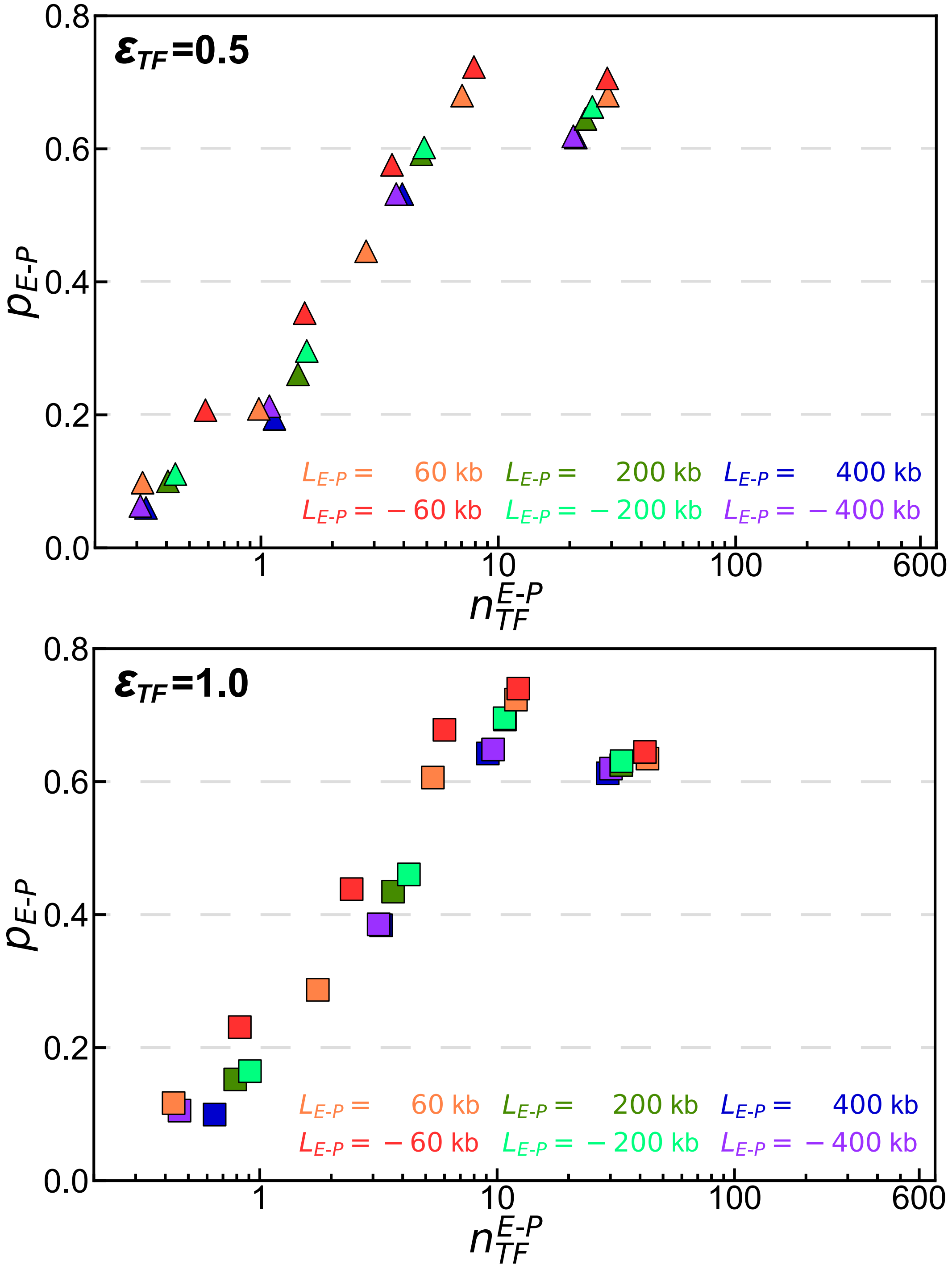


**Figure S2:** Quantification of the dependence of $p_{E\text{-}P}$ on the size of E-P-mediated TF condensates. This analysis is similar to Fig. 2 in the main text, but for $\epsilon_{\mathrm{TF}} = 0.5$ and 1.0, without fitting.

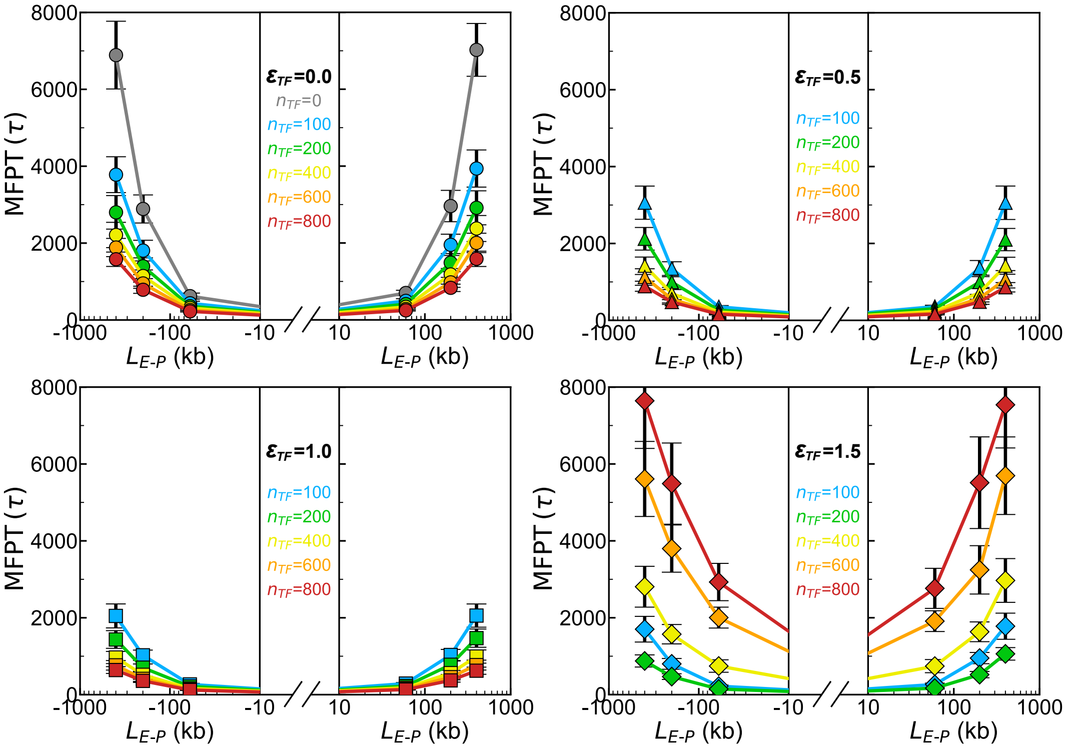


**Figure S3:** MFPT of E-P contact formation. These results are similar to those in Fig. 3 in the main text, but with a linear vertical axis instead of a logarithmic one. The trends of MFPT with respect to E-P genomic length clearly show a reduction in steepness as $\epsilon_{\rm TF}$ or $n_{\rm TF}$ increases.

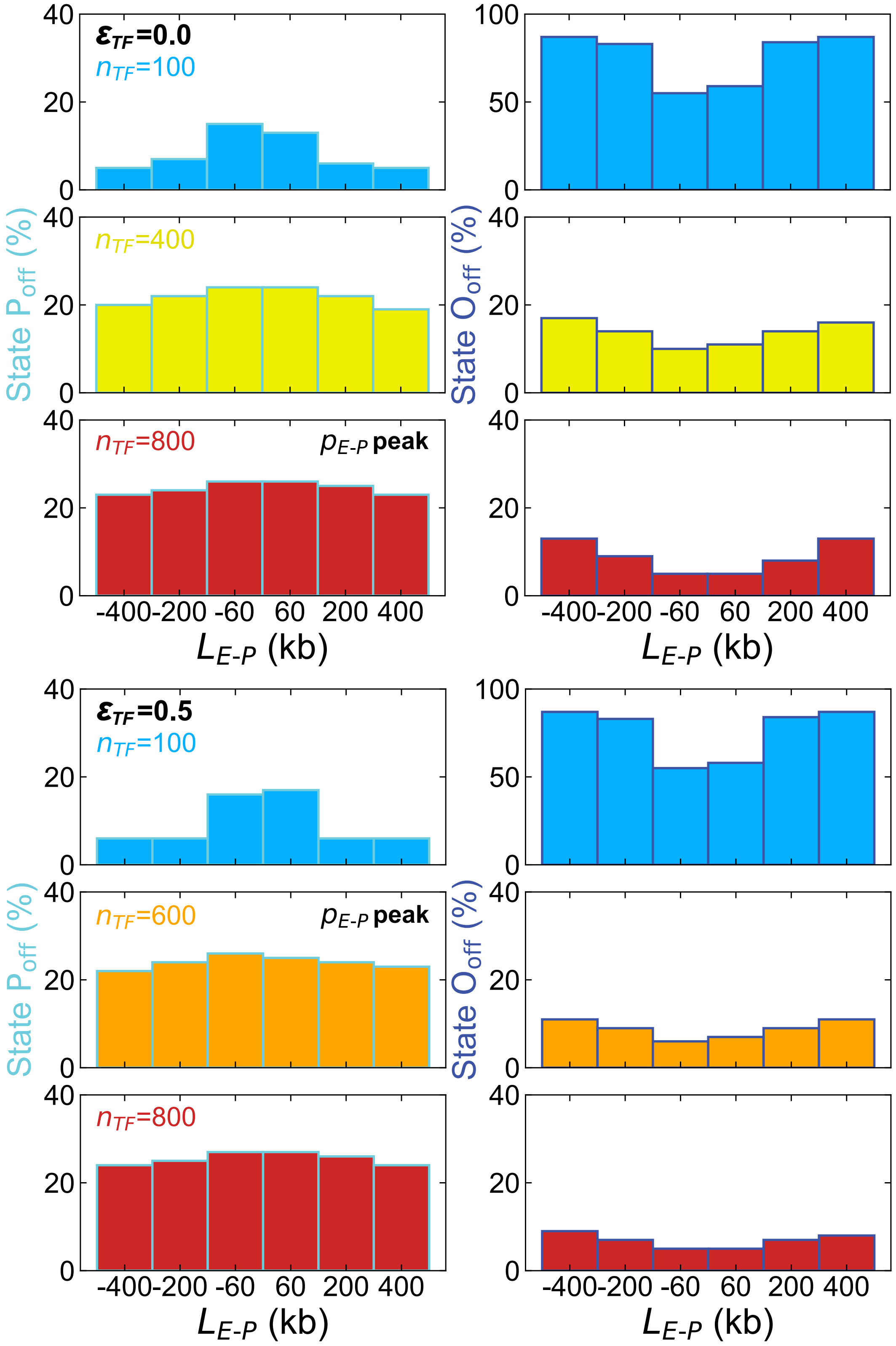


**Figure S4:** Percentage of the $P_{off}$ and $O_{off}$ states for E-P contact. These results are similar to those in Fig. 5 in the main text, but at $\epsilon_{TF} = 0.0, 0.5$.

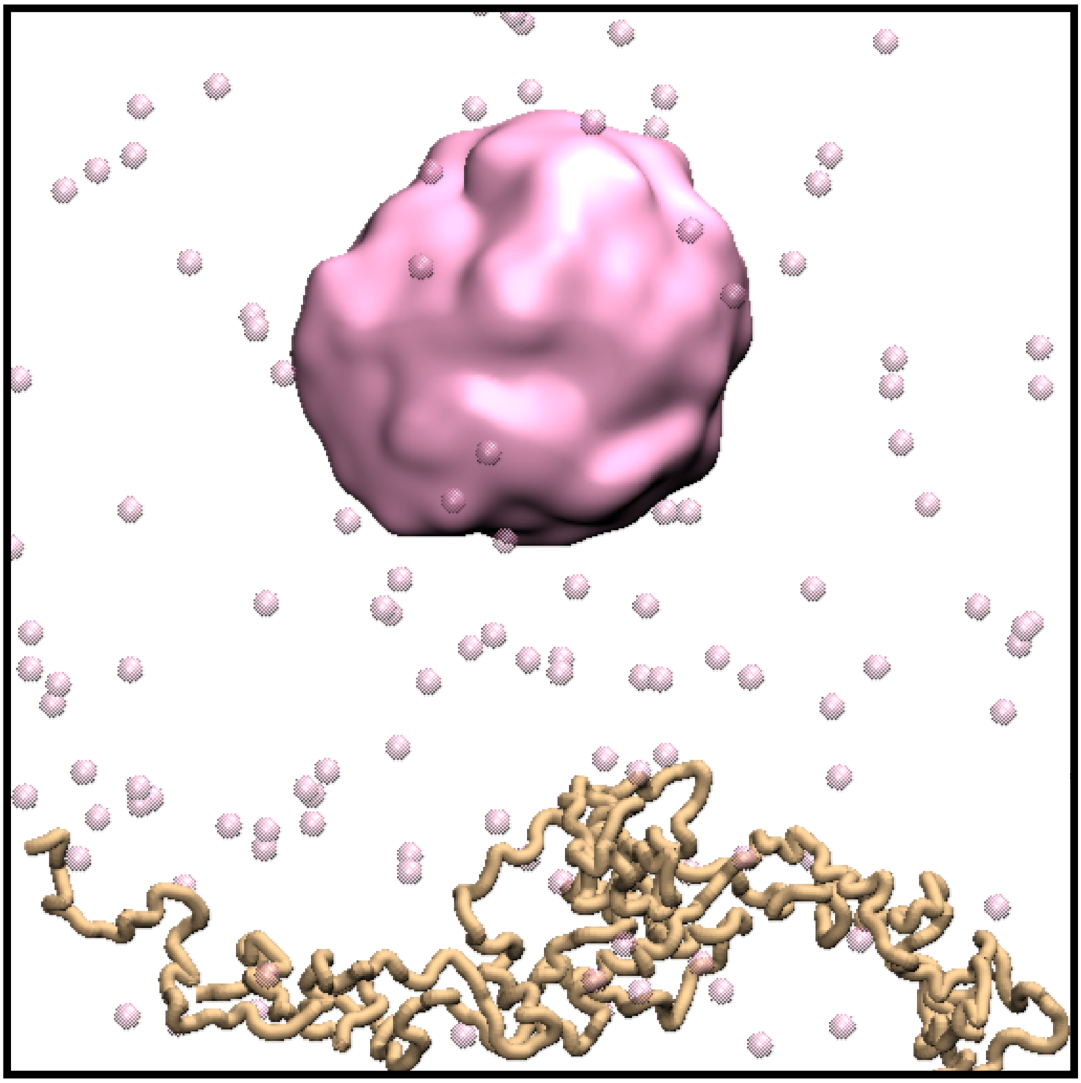

**Figure S5:** Representative snapshot illustrating the self-condensation of TFs without the mediation of E-P loci under conditions of $\epsilon_{\mathrm{TF}} = 1.5$ and $n_{\mathrm{TF}} = 800$. Similar phenomena were also observed when the number of TFs is large and the TF-TF binding affinity is strong (e.g., $\epsilon_{\mathrm{TF}} = 1.5$ and $n_{\mathrm{TF}} = 400, 600$).

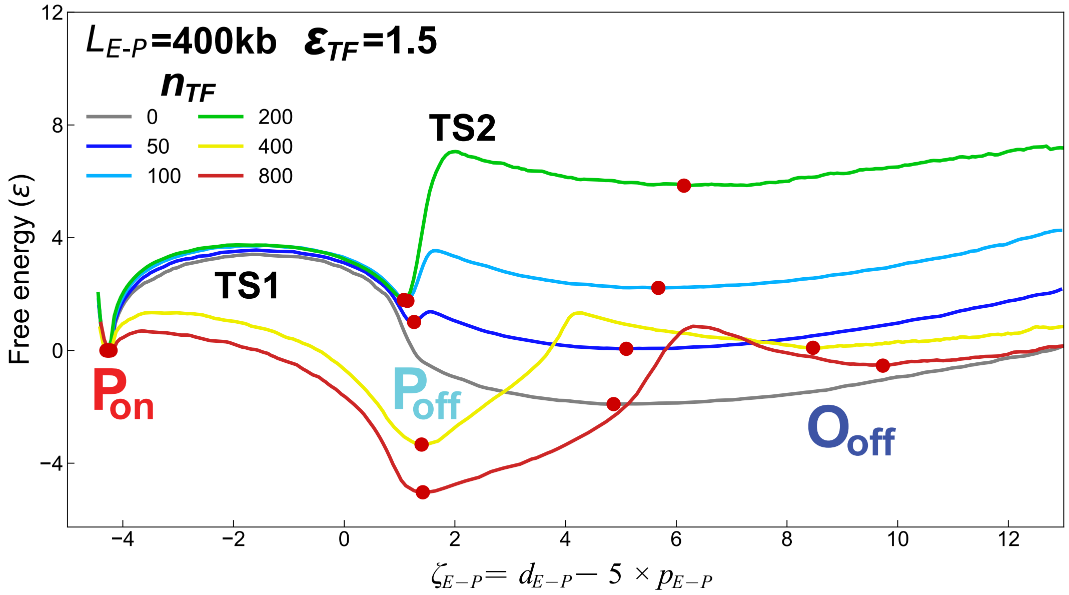


**Figure S6:** 1D free energy landscape projected along the E–P reaction coordinate $\zeta_{E\text{-}P}$ at $L_{E\text{-}P}$ = 400 kb. Distinct stable states ($P_{on}$, $P_{off}$, $O_{off}$) are marked as red points at the free energy minima.

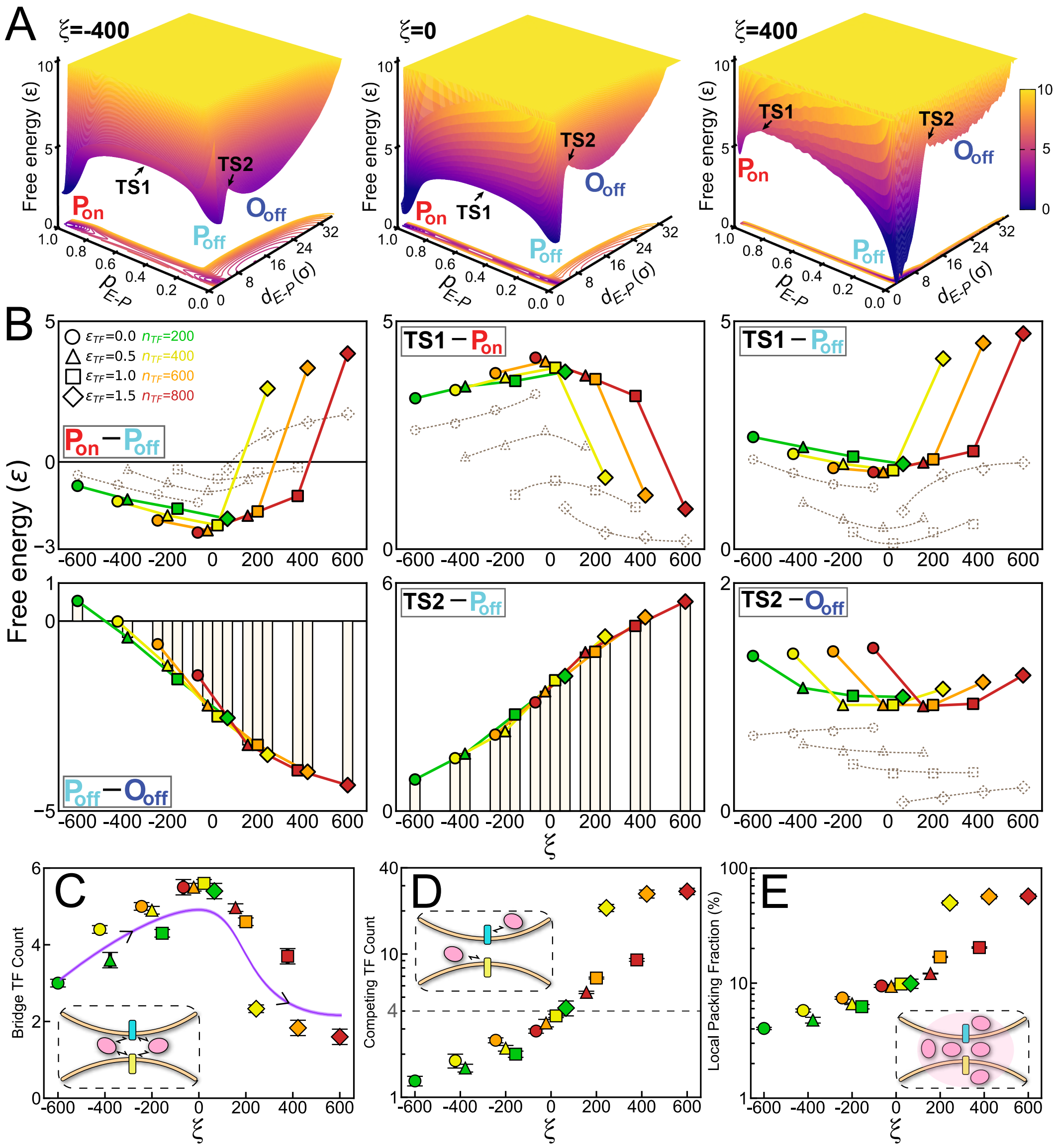

**Figure S7:** Thermodynamic analysis and molecular-mechanistic quantification of E-P contact mediated by TF condensation intensity. These results are similar to those in Fig. 4 in the main text, but at $L_{E\text{-}P} = 200$ kb.

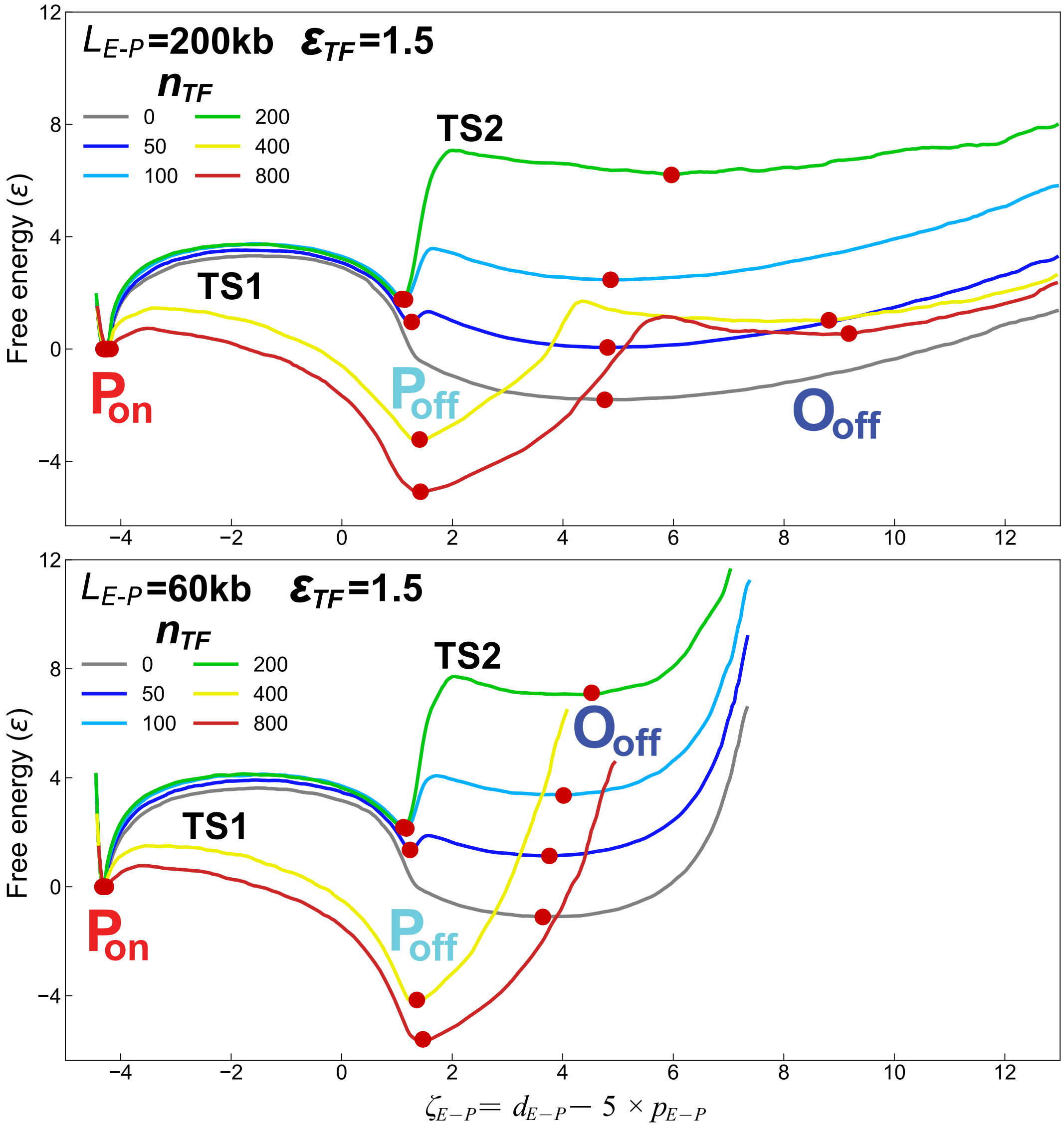


**Figure S8:** 1D free energy landscape similar with Fig. S6 but for $L_{E\text{-}P}$ = 200 and 60 kb. The spatially distant $O_{\text{off}}$ state vanishes under strong TF condensation at $L_{E\text{-}P}$ = 60 kb.
